# Genome-resolved analyses show an extensive diversification in key aerobic hydrocarbon-degrading enzymes across bacteria and archaea

**DOI:** 10.1101/2022.06.22.496985

**Authors:** Maryam Rezaei Somee, Mohammad Ali Amoozegar, Seyed Mohammad Mehdi Dastgheib, Mahmoud Shavandi, Leila Ghanbari Maman, Stefan Bertilsson, Maliheh Mehrshad

## Abstract

Hydrocarbons (HCs) are organic compounds composed solely of carbon and hydrogen. They mainly accumulate in oil reservoirs, but aromatic HCs can also have other sources and are widely distributed in the biosphere. Our perception of pathways for biotic degradation of major HCs and genetic information of key enzymes in these bioconversion processes have mainly been based on cultured microbes and are biased by uneven taxonomic representation. Here we use Annotree to provide a gene-centric view of aerobic degradation of aliphatic and aromatic HCs in a total of 23446 genomes from 123 bacterial and 14 archaeal phyla. Apart from the widespread genetic potential for HC degradation in *Proteobacteria, Actinobacteriota, Bacteroidota*, and *Firmicutes*, genomes from an additional 18 bacterial and 3 archaeal phyla also hosted key HC degrading enzymes. Among these, such degradation potential has not been previously reported for representatives in the phyla UBA8248, Tectomicrobia, SAR324, and Eremiobacterota. While genomes containing full pathways for complete degradation of HCs were only detected in *Proteobacteria* and *Actinobacteriota*, other lineages capable of mediating such key steps could partner with representatives with truncated HC degradation pathways and collaboratively drive the process. Phylogeny reconstruction shows that the reservoir of key aerobic hydrocarbon-degrading enzymes in Bacteria and Archaea undergoes extensive diversification via gene duplication and horizontal gene transfer. This diversification could potentially enable microbes to rapidly adapt to novel and manufactured HCs that reach the environment.

## Introduction

According to the biogenic (organic) theory, petroleum hydrocarbons originate from ancient remains of detrital matter buried and diagenetically modified in marine or freshwater sediments. This organic matter is then gradually converted to petroleum compounds enriched in aromatic and aliphatic hydrocarbons (HCs) via the sequential activity of aerobic and anaerobic microorganisms [1][2][3]. In addition to their role in the formation of oil HCs, microbes play a crucial role in the biological integration of these HCs into the actively cycled carbon pool [4]. Microbial HC degradation occurs through a cascade of enzymatic reactions in three main steps: (i) activation and attack of the HC-bond, producing signature intermediate compounds, (ii) conversion of signature degradation intermediates to central cell metabolites, followed by (iii) mineralization to CO2. Microorganisms must overcome and break the stability and energy in carbon-hydrogen bonds in order to degrade HCs. Since HCs are structurally diverse, a plethora of enzymes are involved in their activation and degradation, and consequently, the energy that needs to be invested in the initial degradation varies. Various microorganisms can degrade different HCs according to their enzymatic repertoire and available energy [5]. Microorganisms have evolved to degrade different HCs under both aerobic and anaerobic conditions. However, biodegradation typically occurs much faster under aerobic conditions, in part due to the availability of thermodynamically favorable electron acceptors that leads to higher energy yield [6], but also because of the action of some HC-degrading enzymes requires oxygen as substrate or cofactor. Similar to all biological pathways, rate-limiting key enzymes drive the main steps of HC degradation.

Under aerobic conditions, oxygenase enzymes initiate the degradation of different aliphatic or aromatic compounds by adding one (mono-oxygenase) or two (di-oxygenase) oxygen molecules. Saturated aliphatic compounds such as alkane and cycloalkane (studied here) are in this process converted to their corresponding carboxylic acid. Catechol/gentisate derivatives are intermediate compounds during aerobic degradation of aromatic mono- and polycyclic HCs. They are then de-aromatized via subsequent meta/ortho cleavage. Intermediate compounds produced during the degradation of aliphatic and aromatic HCs converge to the B-oxidation and tricarboxylic acid (TCA) cycle [7]. While enzymes involved in the downstream part of the degradation process are widespread across living cells shared by many metabolic pathways, the mono/di-oxygenase enzymes catalyzing the first hydroxylation of aliphatic/aromatic compounds are crucial for the initial step in the HC degradation process and likely rate-limiting. Accordingly, microorganisms carrying the enzymes for such initial degradation will be rate-controlling drivers of HC degradation.

The capacity of microbial isolates to metabolically degrade oil HCs have been frequently studied [8-11]. However, our knowledge has until recently been mainly limited to cultivated microorganisms. The present study provides a systemic and genome-resolved view of hydrocarbon degradation capability in the growing database of archaeal and bacterial genomes. To provide this extensive view, we compiled a database of enzymes involved in the aerobic degradation pathway of aliphatic and aromatic HCs (toluene, phenol, xylene, benzene, biphenyl, naphthalene). We then explored the distribution of these enzymes in 24692 publicly available archaeal (n=1246) and bacterial (n=23446) genomes via AnnTree [12] and manually confirmed all annotations. We focused on the microbial genomes containing enzymes for complete/near complete degradation of specific HCs and suggest that lineages with the great genetic potential to degrade a broad range of HC compounds can be exploited for bioremediation purposes. We also reconstructed the phylogenetic relationships of the recovered key HC degradation enzymes to investigate their evolution and explore the potential role of horizontal gene transfer. Several microorganisms contain multiple copies of key HC degrading genes across their genome. We thus explored whether these copies are likely to have been acquired through HGT or if they are likely to be paralogs. Having a genome-resolved view we also studied ecological strategies of these microbes to see whether all critical HC degraders adopt similar growth strategy in terms of the canonical r and k-strategistS.

## Results and discussion

### HC degradation across domain Bacteria

alkB/M and almA/ladA genes are alkane mono-oxygenases that initiate the degradation of short (C5-C15) and long-chain alkanes (>C15), respectively. The alkB/M is rubredoxin-dependent, while almA and ladA are flavin-dependent mono-oxygenases. The genes pheA (phenol), xylM (xylene), xylX, todC1, and tmoA (benzene/toluene) for monoaromatic and bphA1(biphenyl), ndoB (naphthalene/phenanthrene) for polyaromatic compounds code for catalytic domains of ring hydroxylating oxygenases (RHOs) that add -OH group(s) to compounds undergoing degradation (**Supplementary Figure S1**). We explored the distribution of these enzymes and associated degradation pathways in a total of 23446 representatives out of 143512 bacterial genomes available in release 89 of the GTDB database that has been annotated via Annotree [12]. These annotated genomes are dominated by representatives of phyla *Proteobacteria* (32.5%), *Actinobacteriota* (13.3%), *Bacteroidota* (12.13%), and *Firmicutes* (8.01%) (**Supplementary Figure S2** and **Supplementary Table S4**). Among the 123 represented bacterial phyla, 58 phyla had ≤ five genomes available per phylum and combined only represented 0.57% of the explored genomes. To avoid misinterpretations due to this uneven taxonomic distribution of representative genomes, we explored the contribution of members of each phylum in the HC degradation process by showing what proportion of microbes containing each HC degrading enzyme exist in each phylum (panel A of **Supplementary Figures S3-S6**). We also analyze the percentage of members of each phylum containing each HC degrading enzyme to ensure that we consider the contributions of underrepresented phyla in the HC degradation (panel B of **Supplementary Figures S3-S6**).

As expected, representatives of the phylum *Proteobacteria* (*Pseudomonadales* and *Burkholderiales* orders) presented the highest abundance of aliphatic and aromatic HC degrading enzymes, followed by *Actinobacteriota* and *Bacteroidota* for aliphatic and *Actinobacteriota* and *Firmicutes* for *aromatic* HC degrading enzymes (**Supplementary Figures S3** and **S4**, panel A).

Underrepresented phyla remain mainly uncultured and are notably underexplored for metabolic potential (58 of 123 phyla, n=131 genomes). Our analyses revealed that representatives of these taxa contain HC degrading enzymes involved in both the initiation and downstream steps of HC degradation processes. For example, phyla *Tectomicrobia* (*Entotheonella*), *Binatota, Firmicutes*_K, and *Firmicutes*_E contained mono-aromatic HC degradation enzymes (**Figure 2**). In addition to these phyla, we annotated enzymes involved in the degradation of aliphatic HC in representatives of phyla SAR324, *Eremiobacterota* (*Baltobacterales*), *Bdellovibrionota*_B, and *Chloroflexota*_B (**Figure 1**).

**Figure 1.**
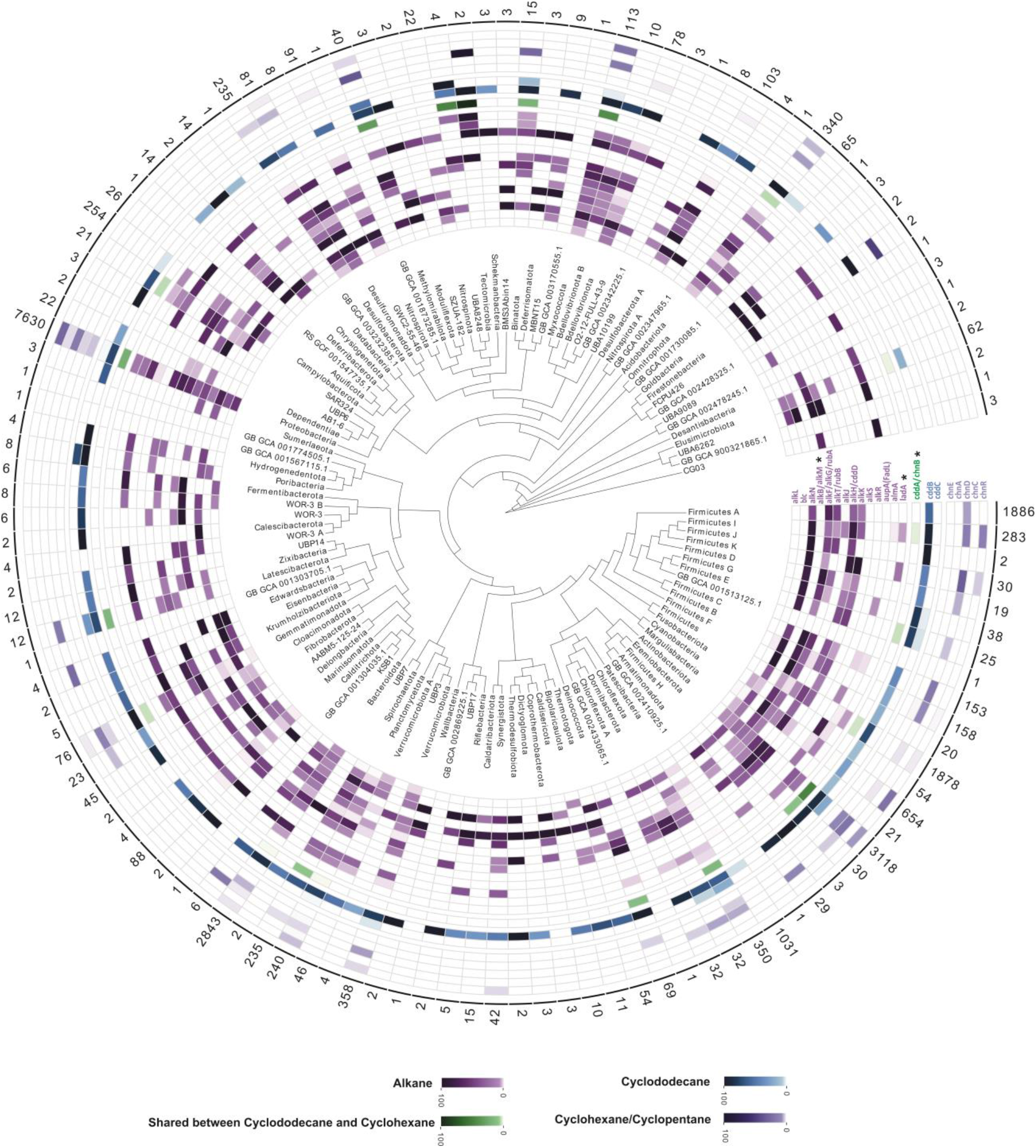
Distribution of aliphatic hydrocarbon-degrading genes across domain bacteria at the phylum level. Each circle of the heatmap represents a gene involved in HC degradation. Various compounds are shown in different colors, as represented in the color legend at the bottom of the figure. Genes marked with an asterisk represent key enzymes of the degradation pathway. Numbers written on each row’s edge indicate the number of screened genomes in that phylum in the AnnoTree website (adopted from GTDB R89). The color gradient for genes of each compound indicates the percentage of HC degrading members of each phylum.

**Figure 2.**
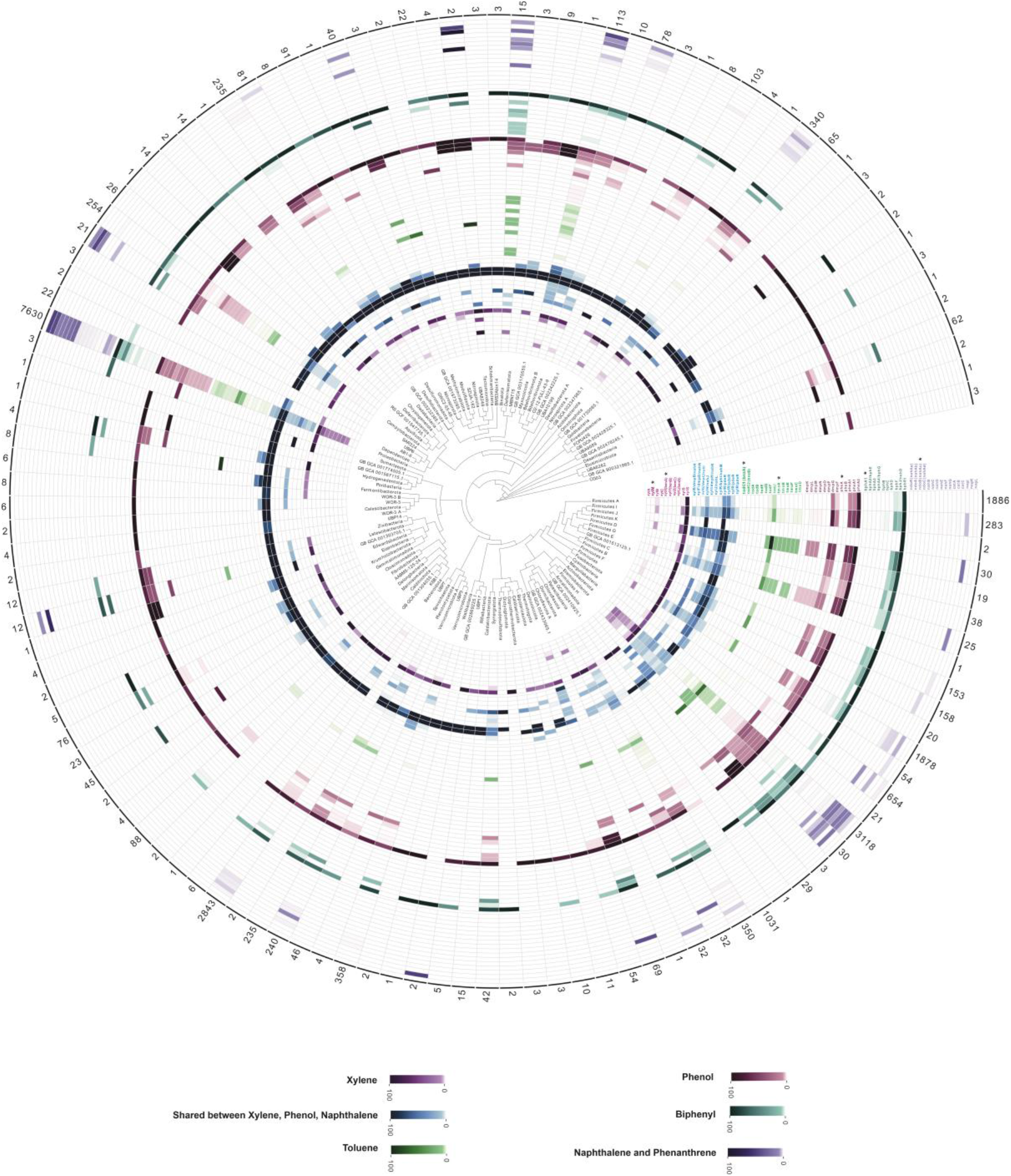
Distribution of aromatic hydrocarbon-degrading genes across domain bacteria at the phylum level. Each circle of the heatmap represents a gene involved in HC degradation. Various compounds are shown in different colors, as represented in the color legend at the bottom of the figure. Genes marked with an asterisk represent key enzymes of the degradation pathway. Numbers written on each row’s edge indicate the number of screened genomes in that phylum in the AnnoTree website (adopted from GTDB R89). The color gradient for genes of each compound indicates the percentage of HC degrading members of each phylum.

Other enzymes in the degradation pathways beyond the key genes for the initial degradation (**Supplementary Table S1**) are typically involved in several degradation pathways and are broadly distributed accordingly. As an example, the process of converting catechol to non-aromatic compounds with further conversion to intermediates of the TCA cycle (e.g., acetaldehyde and pyruvate) (**Supplementary Figure S1**) is shared among degradation pathways of xylene, naphthalene/phenanthrene, and phenol (blue color in **Figure 2**). These ring-cleavage enzymes are also involved in the degradation of aromatic amino acids. Our analysis showed that representatives of phyla Firmicutes (mainly from the orders *Bacillales* and *Staphylococcales*), *Firmicutes*_I, *Firmicutes*_K, *Firmicutes*_E, *Firmicutes*_G, *Firmicutes*_H, *Eremiobacterota, Deinococcota, Chloroflexota, Campylobacterota, Myxococcota* and *Bdellovibrionota* play a significant role in this part of HC degradation process (blue color in **Figure 2**).

### Distribution of key genes involved in the degradation of Alkanes

At lower taxonomic rank, the alkB/M and ladA genes were differently distributed across members of phyla *Gammaproteobacteria, Alphaproteobacteria*, and *Actinobacteriota*, hinting at their capacity for degrading hydrocarbons of variable chain length. Altogether 2089 genomes in orders *Mycobacteriales* (23.95%), *Rhodobacterales* (20.46%), *Pseudomonadales* (17.13%), *Flavobacteriales* (8.3%), *Burkholderiales* (6.16%), *Cytophagales* (3.66%), *Propionibacteriales* (2.47%), *Rhizobiales* (1.89%), and *Chitinophagales* (1.81%) contained alkB/M genes, while ladA was present in 2154 genomes from *Pseudomonadales* (21.05%), *Rhizobiales* (16.27%), *Burkholderiales* (14.44%), *Actinomycetales* (13.44%), *Mycobacteriales* (13.05%), *Bacillales* (4.74%), *Enterobacterales* (3.7%), *Acetobacterales* (2.31%), *Streptomycetales* (1.91%) (**Figures 3** and **4**, panel B, **Supplementary Table S6**).

**Figure 3.**
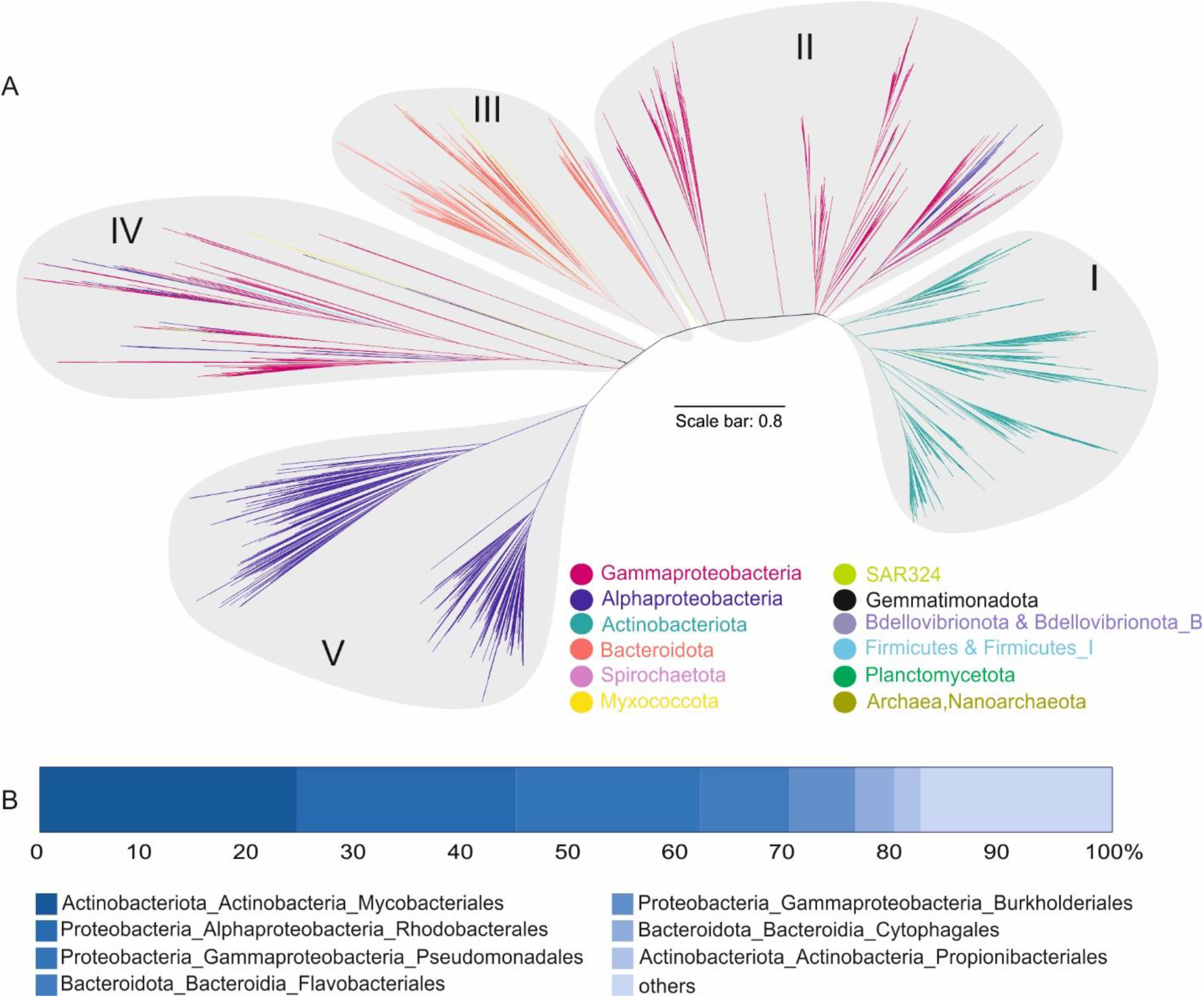
Maximum-likelihood phylogenetic reconstruction of amino acid sequences of alkB/M protein recovered from genomes (short-chain length alkane monooxygenase). A: Major clusters of alkB/M genes according to the reconstructed phylogeny. The scale bar indicates 0.1 branch distance. B: Bar plot representations of the distribution of recovered genes at the order level. The detailed information of the fraction “others” is provided in Supplementary Table S6.

**Figure 4.**
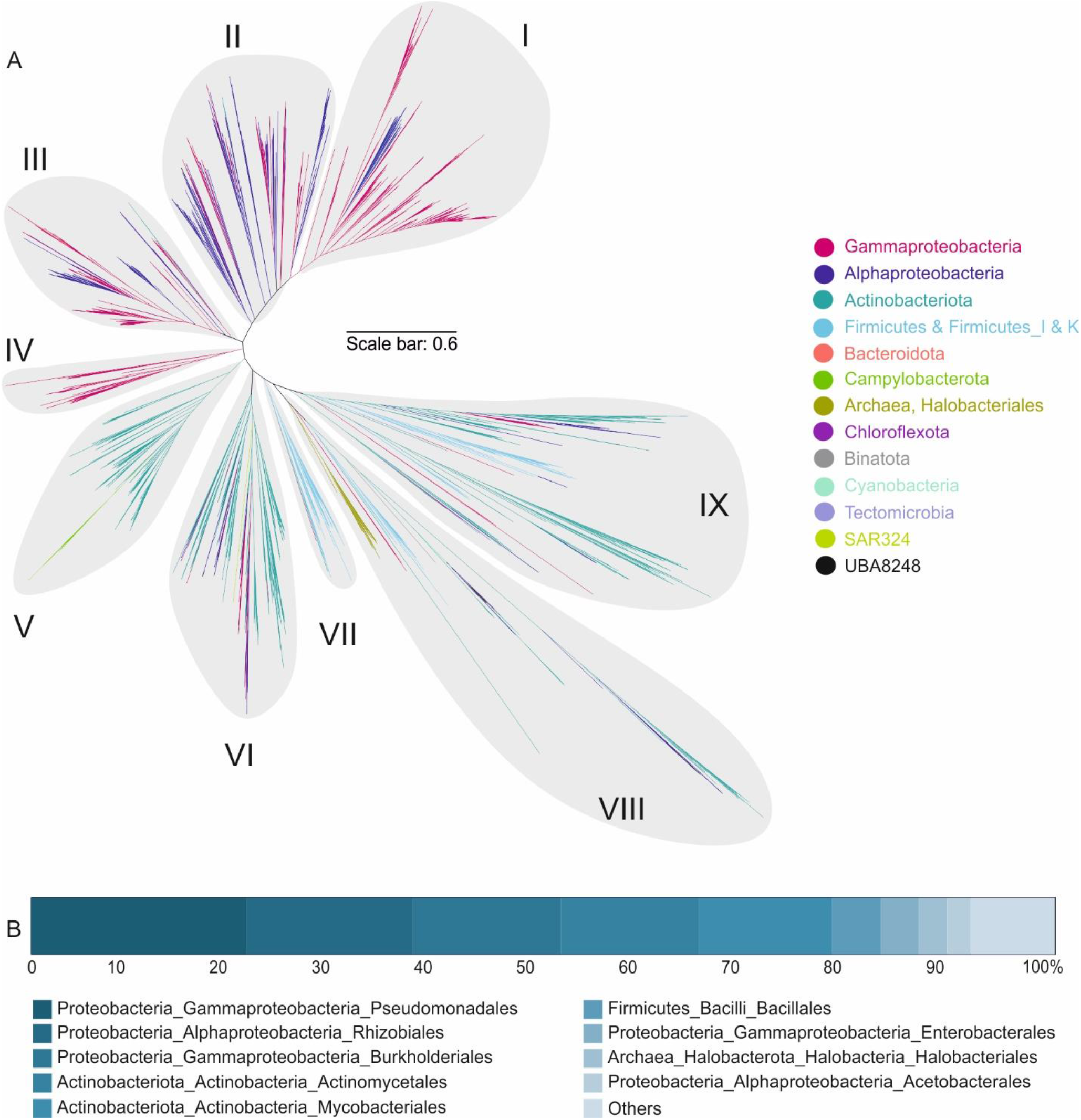
Maximum-likelihood phylogenetic reconstruction of amino acid sequences of ladA protein recovered from genomes (long-chain length alkane monooxygenase). A: Major clusters of ladA genes. The scale bar indicates 0.6 branch distance. B: Bar plot representations of the distribution of recovered genes at the order level. The detailed information of the fraction “others” is provided in Supplementary Table S6.

An indirect role of *Cyanobacteria* in HC degradation, especially in microbial mats, has been previously reported. These primary producers often have the nitrogen-fixing ability and can fuel and promote aerobic and anaerobic sulfate/nitrate-reducing HC degrading microorganisms in microbial mats [13]. There are also reports of a minor role of some *Cyanobacteria* members like *Phormidium, Nostoc, Aphanothece, Synechocystis, Anabaena, Oscillatoria, Plectonema*, and *Aphanocapsa* in direct HC degradation [14][15]. In this study, we detected the presence of long-chain alkane degrading genes, ladA, in different members of *Cyanobacteria* with 0.31 and 12.54% of genomes in this phylum containing ladA (in *Elainella saxicola, Nodosilinea sp000763385*) and almA genes (in *Synechococcales, Cyanobacteriales, Elainellales, Phormidesmiales, Thermosynechococcales, Gloeobacterales, Obscuribacterales*), respectively.

Phylogenetic reconstruction of recovered alkB/M and ladA genes grouped them into five and nine main clades, respectively (**Figures 3** and **4**, panel A). The branching pattern of these clades partially followed the taxonomic signal of the genomes they were retrieved from, specifically for most dominant phyla; however, some branches also contained alkB/M and ladA genes originating from different and distantly related phyla. The placement of phylogenetically diverse groups in one branch is likely to result from the horizontal transfer of these genes between microbial taxa [16]. Additionally, apart from the chromosomal type, both alkB/M and ladA genes have previously been reported to be located on plasmids (OCT and pLW1071), corroborating their potential for horizontal transfer. For instance, there are reports on the intraspecies transfer of alkB/M among *Pseudomonas* members [17]. Placement of rare microbial groups harboring ladA gene among clusters V-IX further suggests a prominent role of Actinobacteriota and Firmicutes members in expanding the distribution of this gene (**Figure 4**).

We also detected several genomes with multiple copies of the alkB gene that were not necessarily branching together in the reconstructed alkB phylogeny, hinting at the probability of either gene duplication, paralogue occurrence or HGT. Examples of these genomes with several copies of alkB/M are *Polycyclovorans algicola* (10), *Nevskia ramose* (7), *Zhongshania aliphaticivorans* (7), *Solimonas aquatic* (7), *Immundisolibacter cernigliae* (6), and *Rhodococcus qingshengii* (6). Multiple copies have also been detected in representatives of the genera Nocardia, Rhodococcus, and Alcanivorax (**Supplementary Table S6**).

Furthermore, the ladA gene was also detected in *Mycolicibacterium dioxanotrophicus, Cryobacterium*_A sp003065485, *Kineococcus rhizosphaerae, Microbacterium* sp003248605, *Paenibacillus*_S sp001956045, *Pararhizobium polonicum, Mycolicibacterium septicum*, and *Microbacterium* sp000799385 with six copies in each genome. Several examples were also present in genera *Pseudomonas*_E, *Bradyrhizobioum, Rhizobioum*, and *Paraburkholderia*, which had more than one copy (904 genomes)(**Supplementary Table S6**).

The presence of multiple copies of alkane hydroxylase genes has been hypothesized to enable cells to use an expanded range of n-alkanes or to adapt to different environmental conditions. However, the exact evolutionary rationale has not yet been established [18][19]. To evaluate this hypothesis, we compared different sequences of each gene in an individual genome (mentioned above for ladA and alkB) using BLAST (**Supplementary Table S7**). The results showed that the identity of multiple gene copies in a single genome was in the range of 30 to 70 percent, while they are still predicted to have the same function. This further supports the hypothesis that these genes originated from different sources and were transferred horizontally.

### Distribution of key genes involved in the degradation of ring hydroxylating oxygenases (RHOs)

Genomes containing RHOs (2761 genomes, 16 phyla) present an overall lower phylogenetic diversity than alkane mono-oxygenases (4669 genomes, 21 phyla for both alkB/M and ladA). In general, alkB/M and ladA enzymes consist of FA_desaturase (PF00487) and Bac_luciferase-like mono-oxygenase (PF00296) domains, respectively (**Supplementary Table S5**). They act non-specifically on a wide range of alkanes of different chain lengths. Therefore, they are likely to be more widespread in genomes, especially because alkane compounds do not exclusively originate from petroleum. For instance, in pristine marine ecosystems, primary producers such as *Cyanobacteria* can release long chain-length aliphatic compounds (e.g., pentadecane, heptadecane). Alkane-producing *Cyanobacteria* include prominent and globally abundant genera such as *Prochlorococcus* and *Synechococcus*. Therefore, marine microorganisms are broadly exposed to aliphatic compounds with different chain lengths, even in environments without oil spills or industrial influence. This can explain why marine ecosystems host a plethora of hydrocarbonoclastic bacteria [20][21].

Enzymes xylX, ndoB, bphA1, and todC1 are composed of two pfam domains, PF00355 (Rieske center) and PF00848 (Ring_hydroxyl_A). These common domains impact the branching in the phylogenetic tree and lead to the neighboring branching of these enzymes (**Figure 5**).

**Figure 5.**
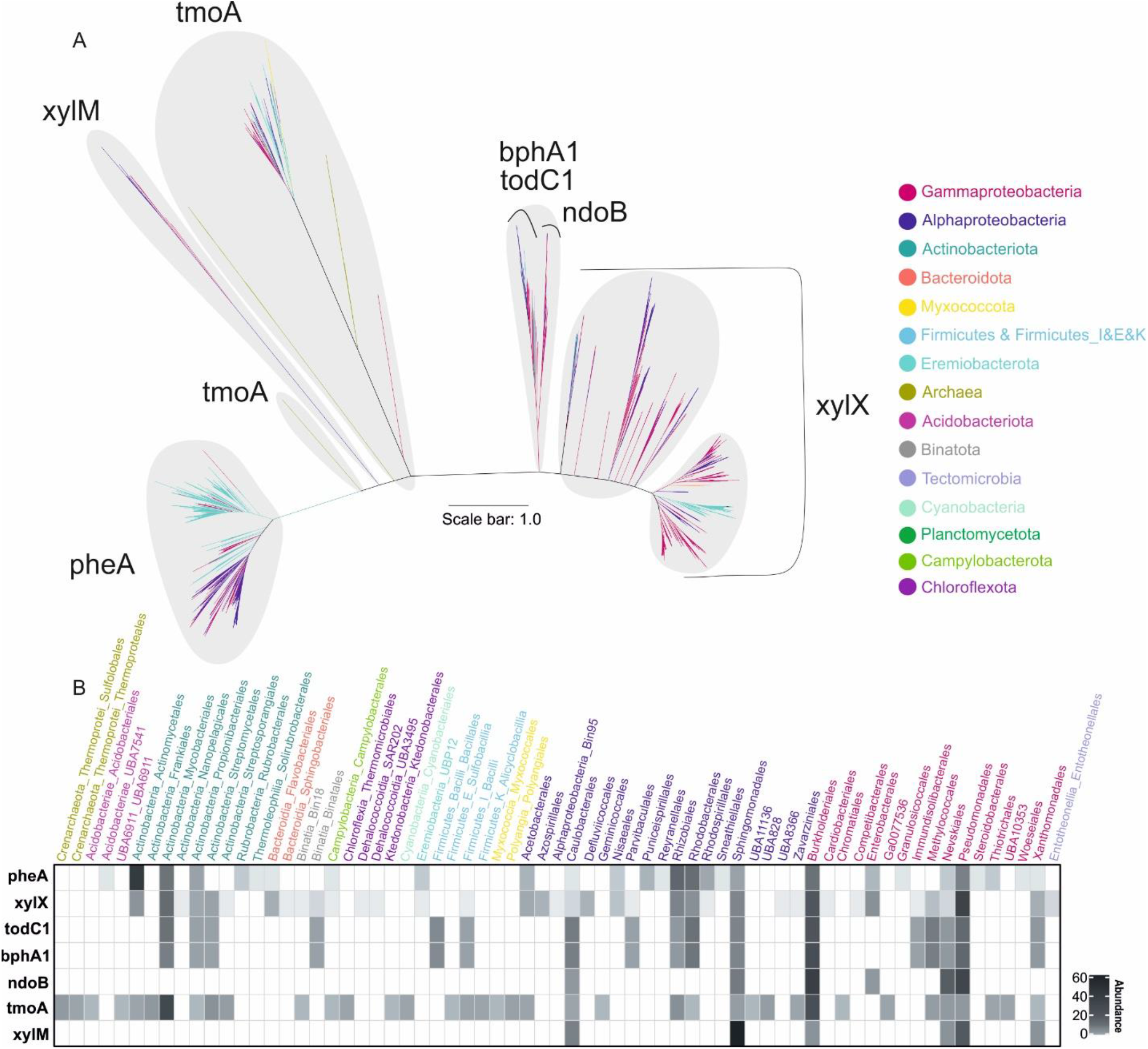
Maximum-likelihood phylogenetic reconstruction of amino acid sequences of ring-hydroxylating oxygenase (RHO) protein recovered from genomes. A: Major clusters of RHO genes. The scale bar indicates 1.0 branch distance. B: Heatmap representations of the distribution of recovered genes at the order level.

RHO enzymes are predominantly present in *Burkholderiales, Pseudomonadales, Sphingomonadales, Caulobacterales*, and *Nevskiales* orders of the phylum *Proteobacteria* (35 different Proteobacterial orders) (Figure 5, B part). However, a significant number of pheA and, to a lesser degree, xylX and tmoA enzymes were also present in Actinobacteriota phylum (9 different *Actinobacteriotal* orders) (**Figure 5**, B part).

*Sphingomonadales* are prominent bacteria in the rhizosphere and are also abundant in littoral zones of inland waters. Accordingly, we suggest that these bacteria may have evolved a capacity to degrade different aromatic compounds in response to the high concentrations of aromatic secondary metabolites typically seen in the plant rhizosphere. Additionally, *Sphingomonadales* are known for their large plasmids with intraspecies transmission [22].

Among all investigated RHO genes, the highest phylogenetic diversity was observed in tmoA (208 genomes in 12 phyla and 38 orders) and xylX (1486 genomes in 9 phyla and 38 orders) genes (**Figure 5**, B part). In the case of tmoA gene, it might be due to the wide range of HC compounds susceptible to this enzyme (e.g., benzenes, some PAHs, and alkenes)[23][24]. Therefore, more diverse genera harbor tmoA gene and can degrade different types of HCs.

Underrepresented microbial groups with a limited number of RHO genes also featured tmoA, xylX, and pheA genes. *Myxococcota, Acidobacteriota, Chloroflexota, Firmicutes*_I,E,K, and *Cyanobacteria* with tmoA gene were clustered separately, reflecting their distinct protein sequence and the lower possibility of HGT among these groups. For xylX, *Eremiobacterota* affiliated genes were placed together with genes from *Gammaproteobacteria*, and *Tectomicrobia, Binatota, Chloroflexota*, and *Firmicutes*_I were placed in separate branches near *Actinobacteriota*. In addition, *Acidobacteriota, Eremiobacterota*, and *Campylobacteria* with pheA gene were nested within *Alphaproteobacteria* members. The phylogeny of RHO genes was also more consistent with taxonomy than the phylogeny of alkB/M and ladA.

*Bionatota*, a recently described phylum shown to be efficient in HC degradation, harbored todC1, bphA1 (in Binatales order), and xylX (Bin18, *Binatales*) genes from RHOs and ladA (in Bin18) from alkane hydroxylases. Representatives of this phyla have been reported to play a role in methane and alkane metabolism [25]. However, we also noted the further potential of Binatales and Bin18 orders of this phylum in aromatic HC degradation.

RHOs can be located either on the chromosome or plasmid, depending on the organism. For instance, todC1, bphA1, and tmoA genes were reported to be on the chromosome [26], while in another study, they were detected on a plasmid [27]. Other RHOs, including xylX, xylM, pheA, and ndoB have mainly been reported to be hosted by plasmids [26][24].

Multiple copies of RHO genes in one genome were detected for xylX and pheA. *Immundisolibacter cernigliae surprisingly* contained 21 variants of xylX. This genome also had six copies of alkB/M and was isolated from a PAH-contaminated site [28]. The high HC degradation potential of other members of this genus has also been reported in the marine ecosystem [29][30]. *Rugosibacter aromaticivorans* (containing 5, 2 and 2 copies of xylX, ndoB, and tmoA genes, respectively), *Pseudoxanthomonas* A spadix B (with 4, 2 and 2 copies of xylX, todC1 and bphA1 genes, respectively), *Thauera* sp002354895 (4), *Pigmentiphaga* sp002188635 (4) are other examples of genomes that have multiple copies of the xylX gene. Although xylX gene was detected in *Actinobacteriota*, multiple copies in a genome were seen only among the *Proteobacteria phylum*.

The BLAST identity among variants of the xylX gene in *Immundisolibacter cernigliae* ranged between 35 to 81 percent. Three sequences of these 21 xylX copies (xylX 18, 19, and 22, in **Supplementary Figure S7**) showed higher BLAST identity with the xylX gene of the *Rugosibacter* genus than other copies in the *Immundisolibacter cernigliae* genome itself (**Supplementary Table S7** and **Supplementary Figure S7**). Several xylX copies of I. *cernigliae* (10, 11, 13, and 15) had more edges than others in the network, and their interactions (**Supplementary Figure S7, highlighted in red**) represent their similarity with xylX copies of *Caballeronia, Sphingobium*, and *Pseudoxanthomonas, Pseudomonas*, and *Thauera* genera. In addition, xylX 5 and 7 of *Immundisolibacter* had almost similar blast identity with *Pigmentiphaga* genus and other xylX copies in I. *cernigliae*. This suggests that multiple copies of the xylX gene in I. *cernigliae* potentially originated from horizontal transfer.

On the other hand, *Glutamicibacter mysorens* (4), *Enteractinococcus helveticum* (4), and many other genomes from the *Castellaniella, Kocuria*, and *Halomonas genera*, had several pheA copies in their individual genomes. To a lesser degree, tmoA gene was present in multiple copies in *Pseudonocardia dioxanivorans* (4), *Rhodococcus* sp003130705 (3), *Amycolatopsis rubida* (3) and *Zavarzinia compransoris*_A (3) genera.

While bphA1 and todC1 have different KO identifiers (**Supplementary Table S1**), our manual checks showed that they had the same conserved domain based on NCBI CD-Search [31]. We kept both annotations for cases where one gene was annotated with both KO identifiers. Previous studies also report similar homology and substrate specificity between todC1 and bphA1 [27].

xylM, as one of the enzymes mediating the initial steps in toluene/xylene degradation, showed the lowest abundance and phylogenetic diversity (27 genomes in 1 phylum and 6 orders). Toluene/benzene can generally be degraded through different routes and three of the most prevalent approaches were studied here. xylX, todC1, and tmoA are the initial oxygenase enzymes of these three pathways. They are diverse in starting the degradation and composed of different domains, while downstream degradation converges to catechol derivatives as intermediates. xylM can also initiate toluene degradation in addition to xylene. xylX then converts produced benzoate to catechol. Therefore, while we report a lower diversity of genomes harboring xylM, there are alternative degradation pathways in different microorganisms that can degrade the same compound.

As the number of rings in aromatic compounds increases, the number and diversity of microbial groups capable of degrading them decreases, and microbial groups with ndoB (naphthalene 1,2-dioxygenase) accordingly showed the lowest abundance after xylM gene. The genomes hosting ndoB had limited phylogenetic diversity (35 genomes in 1 phylum and 6 orders) and were found mainly in representatives of *Alphaproteobacteria* (*Sphingomonadales* (17) and *Caulobacterales* (2)) and *Gammaproteobacteria* (*Pseudomonadales* (5), *Burkholderiales* (1), *Nevskiales* (1)).

### Ecological strategy of HC degrading bacteria

Microorganisms are broadly divided into two main functional growth categories, i.e., oligotrophic/slow-growing/k-strategist or copiotrophic/fast-growing/r-strategist. These ecological strategies are associated with the genome size that, in turn, directly correlates with the GC content [32]. To get further insights into the ecological strategies of organisms that feature HC degrading genes, we compared the distribution of GC content and estimated genome size. This analysis revealed that HC degrading genes were present in genomes with a broad genome size range (1.34 to 16.9 Mb) and GC content (26.9 to 76.6 %) (**Supplementary Figure S8**, data available in **Supplementary Table S8**). Genomes with GC percent equal to or lower than 30% mainly had alkB gene and belonged to representatives of the Flavobacteriales order (genome sizes in the range of 1.4 to 4.2 Mb). The largest genome studied here, *Minicystis rosea* from the phylum Myxococcota (genomes size of 16.9 Mb), also contained alkB. The alkB gene of *Minicystis rosea* phylogenetically clustered together with homologs from Gammaproteobacteria representatives (*Immundisolibacter* and *Cycloclasticus genera*) (**Figure 3**). The large genome size of *Minicystis rosea* and its alkB gene placement together with the Gammaproteobacteria in the reconstructed phylogeny suggests horizontal transfer for this gene to *Minicystis rosea*. These analyses suggest that HC degradation ability is present in both k-strategist and r-strategists microorganisms. Earlier studies have shown that r-strategist serves as the principal HC degraders after oil spills and at other point sources of pollution in marine environments [33-36]. Indeed, most obligate hydrocarbonoclastic bacteria are r-strategists (Proteobacteria domain) and are mainly reported to be isolated from marine samples [37]. This group is adapted to live in oligotrophic environments with transient nutrient inputs and rapid consumption of substrates upon episodic inputs by means of fast growth and population expansion [38]. In contrast, reports of oil-polluted soil samples suggest a predominance of k-strategists, especially in the harsh conditions (High concentration of HC, soil dryness, etc.) commonly seen in many such soil environments [39-41]. Hosting multiple copies of genes coding for HC degrading enzymes seems to be a shared feature in both r-and k-strategists with small and large genome sizes alike and appears to be a universal evolutionary strategy for HC degradation.

### Genome-level analysis of HC degradation

Microorganisms are known to use division of labor or mutualistic interactions to perform HC degradation in the environment [42][43]. However, 92 genomes (less than 0.5%) of 23446 investigated bacterial genomes do in fact contain all the enzymes required to degrade at least one HC compound completely. These 92 genomes all belong to *Actinobacteriota* (n=25) and *Proteobacteria* (n=67)(Figure 6).

**Figure 6.**
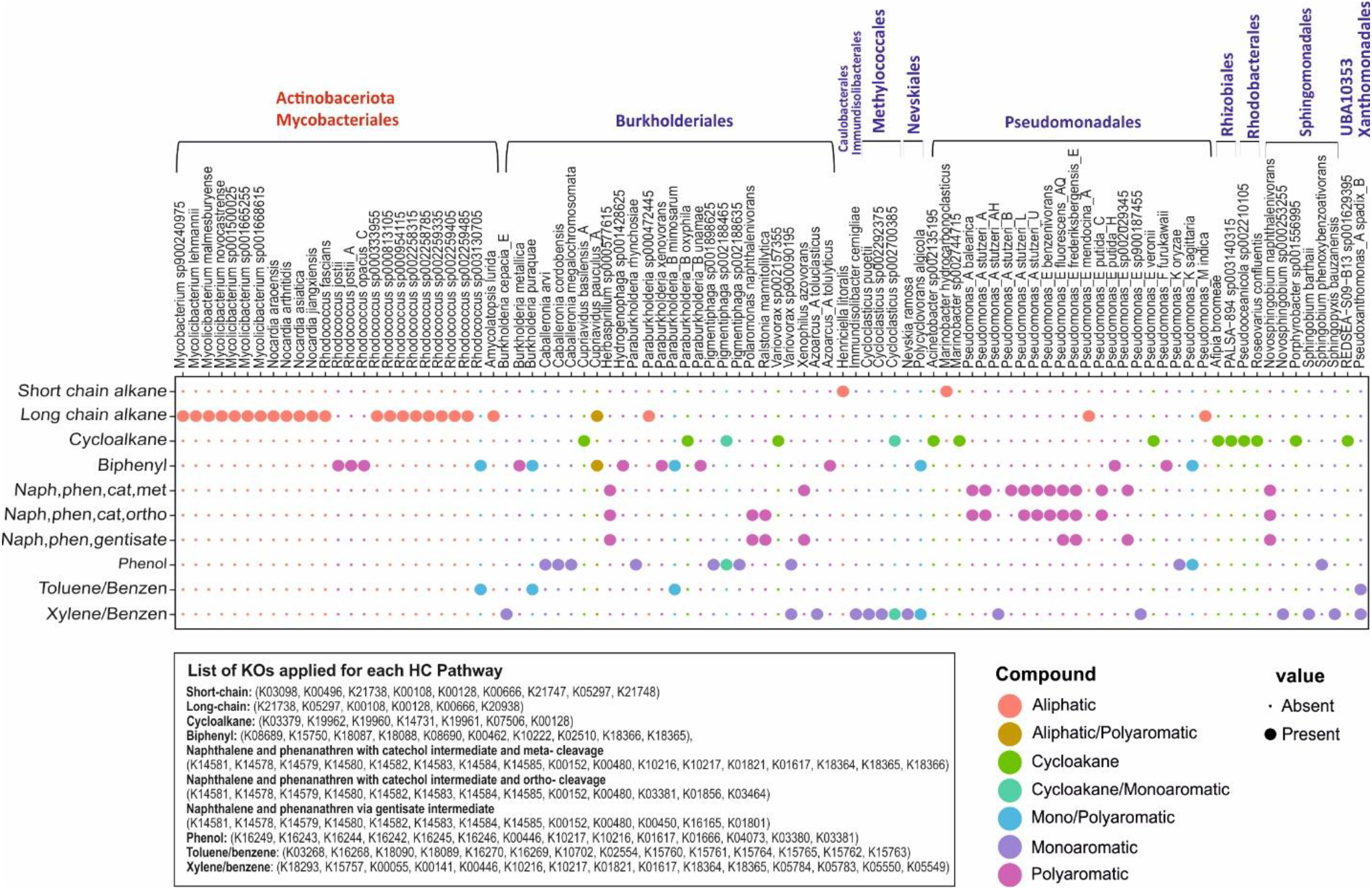
Genomes with complete/near complete degradation pathways of different HCs. Colors represent the type of HC that microbial genomes could degrade. Rows represent the type of HCs and columns show the name of genomes. Orders belonging to Proteobacteria and Actinobacteriota phyla are written in blue and red, respectively. KEGG orthologous accession number of enzymes for the complete degradation process of each compound is written at the figure’s bottom.

Microorganisms have evolved two pathways for naphthalene degradation that involve the production of either catechol or gentisate as aromatic degradation intermediate (**Supplementary Figure S1**). Catechol can in turn, be further degraded via meta-or -ortho cleavage. Several microorganisms, including *Novosphingobium naphthalenivorans, Pseudomonas*_E *fluorescens*_AQ, *Pseudomonas*_E *frederiksbergensis*_E, and *Herbaspirillum* sp000577615, feature both of the mentioned pathways and even have the ability to perform ortho and meta cleavage simultaneously (**Figure 6**).

Moreover, *Cupriavidus pauculus*_A (long-chain alkanes and also biphenyl), *Cycloclasticus* sp002700385 and *Paraburkholderia*_B *oxyphila* (*Cycloalkane* and xylene/benzene), *Pigmentiphaga* sp002188465 (Cycloalkane and phenol), *Rhodococcus* sp003130705, *Burkholderia puraquae*, and *Paraburkholderia*_B *mimosarum* (Toluene and biphenyl) can degrade more than one HC compound autonomously (**Figure 6**). Members of Burkholderiales were able to degrade even more diverse compounds individually, while *Actinobacteriota* representatives mainly contribute to the degradation of aliphatic compounds. This ability was also apparent in **Figures 1, 3**, and **4**. The potential for autonomous HC degradation wasn*’*t detected in genomes of more rare bacterial phyla. Moreover, none of the archaeal genomes investigated in this study contained all genes for the complete degradation of HCs.

### HC degradation across domain archaea

Generally, HC degradation ability seems to be less prevalent among archaea as compared to bacteria. The phylum *Halobacterota* had the highest proportion of enzymes involved in the degradation of both aliphatic (n=14 enzymes of aliphatic degradation pathway) and aromatic (n=25 enzymes of aromatic degradation pathway) compounds among the studied archaea (**Supplementary Figure S9**). The alkB enzyme, responsible for short-chain alkane degradation, was detected in two copies in a single member of the phylum Nanoarchaeota (ARS21 sp002686215). This gene was clustered together with alkB identified in Gammaproteobacteria representatives (GCA-002705445 order) (**Figure 3**). Genes needed to initiate degradation of long-chain alkanes and cyclododecane/cyclohexane as well as cyclopentane degradation via ladA and cddA/chnB genes were more prevalent among *Halobacterota* representatives (75 genomes in 7 families; *Haloferacaceae, Haloarculaceae, Natrialbaceae, Halococcaceae, Halalkalicoccaceae, Haloadaptaceae*, and *Halobacteriaceae*) (**Figures 4** and **Supplementary Figure S9**). Among investigated RHOs, only tmoA that initiates toluene degradation was present in 5 Sufolobales and 2 Thermoproteales genomes of the phylum Crenarchaeota (**Figure 5**). Detected archaeal tmoA and ladA genes branched separately from bacteria in the phylogenetic trees (**Figures 4 and 5**). Apart from alkB, gene duplications were present in several genomes for both tmoA (*Sulfolobus* and *Acidianus* genera) and ladA (*Halopenitus persicus* and *Halopenitus malekzadehii*).

Key enzymes needed to initiate HC degradation were rarely present in archaea (Figures 3, 4, and 5), indicating that Archaea might not play a significant role in the typically rate-limiting initial degradation of HCs. However, several studies report the ability of halophilic archaeal isolates (e.g., *Halorubrum* sp., *Halobacterium* sp., *Haloferax* sp., *Haloarcula* sp.) to degrade both aliphatic (n-alkanes with chain lengths up to C18 and longer) and aromatic (e.g., naphthalene, phenanthrene, benzene, toluene and *p*-hydroxybenzoic acid) HCs and use them as their sole source of carbon [44-46]. This may imply that archaea carry alternative and hitherto unknown enzymes for triggering HC degradation. However, there is no complete genome information available for the mentioned isolates to screen them for the presence of alternative degrading enzymes [11]. The *Haloferax* sp., capable of using a wide range of HCs as its sole source of carbon, present in the AnnoTree database (RS_GCF_000025685.1), contained none of the key degrading genes. The AnnoTree website chooses representative genomes having completeness of higher than 90%, whcih reduces the likelihood of incompleteness of the studied genome as a reason for the absence of these genes. Therefore, alternative HC degrading genes that are present in the accessory part of the genomes might be responsible for the observed degradation.

On the other hand, the recent reconstruction of three metagenome-assembled Thermoplasmatota genomes (Poseidonia, MGIIa-L2, MGIIb-N1) from oil-exposed marine water samples (not included in the GTDB release89) contained enzymes involved in alkane (alkB) and xylene (xylM) degradation [30]. Hence as these global genome depositories continue to expand, we may have to revise or update our findings.

A total number of 597 archaeal genomes contain enzymes involved in the degradation of aromatic compounds regarding the conversion of catechol to TCA intermediates. This is observed in the phyla *Halobacterota* (176 genomes in *Haloferacaceae, Haloarculaceae, Natrialbaceae, Halococcaceae*,

*Halobacteriaceae, Methanocullaceae, Methanoregulaceae, Methanosarcinaceae, Archaeoglobaceae*, and some other methano-prefixed families), *Thermoplasmatota* (175 genomes in *Poseidoniales*, Marine Group III, *Methanomassiliicoccales*, UBA10834, *Acidiprofundales*, DHVEG-1, UBA9212), and *Crenarchaeota* (110 genomes in *Nitrospherales, Desulfurococcales, Sufolobales, Thermoproteales*). This widespread capacity for degrading downstream intermediates in aromatic HC degradation implies that archaea interact closely with bacteria in HC degradation.

## Conclusions

HCs are ubiquitously distributed in the biosphere and do not exclusively originate from oil. In this study, the distribution of key HC degrading enzymes involved in the degradation of certain HCs (aliphatic and aromatic types) is provided at genome resolution for both the archaeal and bacterial domains. Extensive environmental genome and metagenome sequencing over the last decades has significantly increased the number of available microbial genomes and enriched contemporary genomic databases. The genome-based taxonomy using average nucleotide identity (ANI) or relative evolutionary divergence adopted by the Genome Taxonomy Database; GTDB [47,48] as a reproducible method has in parallel revised and updated some taxonomic ranks. The order Oceanospirillales, as an example, is a well-known taxon in the marine oil degradation context, and its representatives have been frequently reported as one of the main HC degrading members in response to oil pollution [49,50,37]. Nonetheless, this taxonomic rank has been removed from the genome-based taxonomy, and its members have been mainly placed in the order Pseudomonadales [51]. This could potentially cause a communication gap between the existing literature and new research. An updated comprehensive metabolic survey of Bacteria and Archaea for HC degradation potential at genome resolution could thus help bridge this gap. Our extensive survey shows that a greater diversity of bacteria is involved in aliphatic HC degradation compared to aromatic HCs. Few genomes were detected to contain all necessary enzymes to carry out complete degradation pathways. This reiterates previous findings that microbes generally cooperate for HC degradation by “division of labor” and a community perspective would therefore be crucial to predicting the fate of oil HCs in the ecosystem. We detected HC degrading ability among both r and k strategists and found signals of gene duplication and horizontal transfer of key HC degrading genes. This could be an efficient way to increase degradation capability among microbial members and potentially help them adapt to the available pool of HCs in their ecosystem.

## Materials and methods

### Data collection of HC Degrading enzymes

Representative compounds from each category of HCs, including saturated aliphatic (short/long-chain alkanes) and alicyclic (cyclohexane/cyclododecane), compounds with mono-aromatic (toluene, phenol, xylene, and benzene), and poly-aromatic (PAHs) (naphthalene, phenanthrene, and biphenyl as representatives) hydrocarbons were selected to survey the distribution of Bacteria and Archaea capable of their degradation under aerobic conditions.

A complete list of enzymes involved in the degradation pathway of mentioned HCs was compiled from previous reports [52-57]. We explored these enzymes in Kyoto Encyclopedia of Genes and Genomes (KEGG)[58], Pfam [59], TIGRFAMs [60], InterPro [61], and UniProt [62] databases. The accession number of enzymes in each mentioned database, their function, name, reaction (if available), EC number, and additional information are provided in **Supplementary Table S1**.

### Distribution of HC degrading enzymes among bacterial and archaeal representative genomes

The distribution of the compiled HC degrading enzymes described in **Supplementary Table S1** was assessed across domains Bacteria and Archaea using AnnoTree (http://annotree.uwaterloo.ca) [12]. AnnoTree database is providing functional annotations for 24692 genome representatives in the genome taxonomy database (GTDB) release 89. The phylogenetic classification of genomes is derived from the GTDB database (release R89). In total, the annotation information for 18, 10, and 90 enzymes involved in the degradation process of alkane, cycloalkane, and aromatic HCs, respectively, were analyzed. Genome hits were collected at the thresholds of percent identity ≥ 50, e-value cut off ≤1e^-5^, subject/query percent alignment ≥70 for KEGG annotations, and e-value cut off ≤ 1e^-5^ for Pfam and TIGRFAMs annotations. For each HC degrading enzyme, we first checked KEGG annotations. If there were no KEGG accession numbers for the enzyme, the second priority was TIGRFAMs; otherwise, the Pfam annotation was considered. The table contains information for the distribution of HC degrading enzymes of each pathway present in representative genomes from bacteria and archaea domains, as is shown in **Supplementary Tables S2** and **S3**, respectively.

### Phylogeny of bacteria and archaea augmented with the abundance of HC degrading enzymes

Evolview, a web-based tool for the phylogenetic tree visualization, management, and annotation, was used to present the distribution view of HC degrading enzymes in representative genomes across bacterial/archaeal phylogenomic trees [63][64].

The phylogenomic tree of bacteria and archaea in the Newick format, at the phylum level (123 and 14 leaves, respectively), was adopted from the AnnoTree website (November 21^st^, 2020). Trees were uploaded as the reference tree in Evolview. According to the abundance tables of HC degrading enzymes prepared for each degradation pathway, four heatmaps were plotted for bacteria and archaea domains (separately for aliphatic and aromatic compounds).

### Single gene phylogeny

To provide the evolutionary history of key enzymes in each HC degradation pathway, the protein sequence of that enzyme was manually confirmed by inspecting their conserved domains using the NCBI web CD-Search tool (https://www.ncbi.nlm.nih.gov/Structure/bwrpsb/bwrpsb.cgi) [31]. Validated amino acid sequences were then aligned using Kalign3 software [65], and their phylogenetic tree was reconstructed using FastTree2 [66].

## Acknowledgments

The computational analysis was performed at the Center for High-Performance Computing, School of Mathematics, Statistics, and Computer Science, University of Tehran.

## Author contributions

M.M. devised the study. M.R.S., L.G.M., and M.M., performed the bioinformatics analysis M.R.S. and M.M. interpreted the data with input from S.M.M.D., S.B., M.A.A., and M.S.. M.R.S. and M.M drafted the manuscript. All authors read and approved the manuscript.

## Supplementary Figures

**Supplementary Figure S1.**
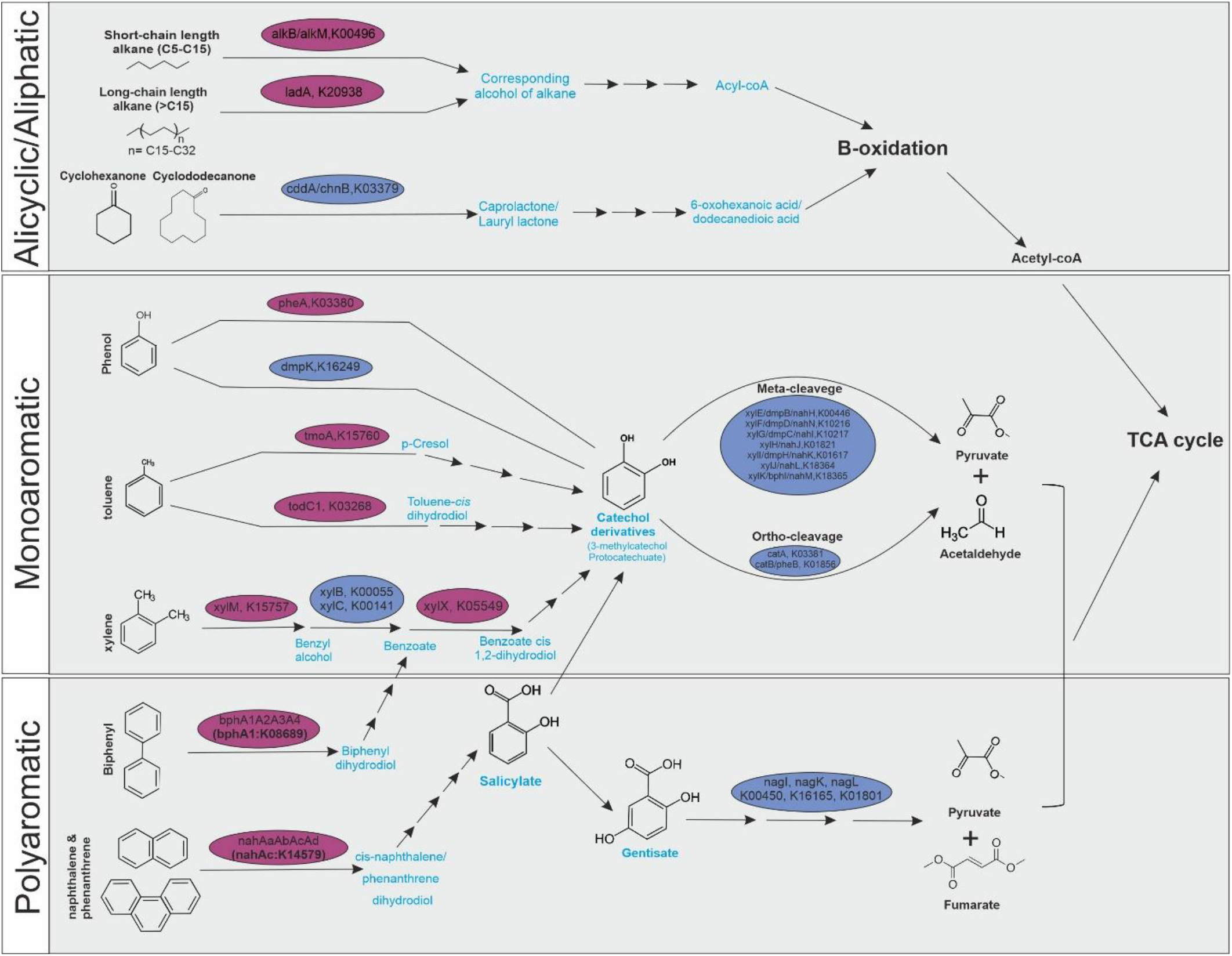
Schematic representation of HC degradation pathways studied in this work. Purple circles show key HC degrading enzymes trigerring the degradation. Blue circles are other crucial enzymes.Important intermediate compounds are written in blue.

**Supplementary Figure S2.**
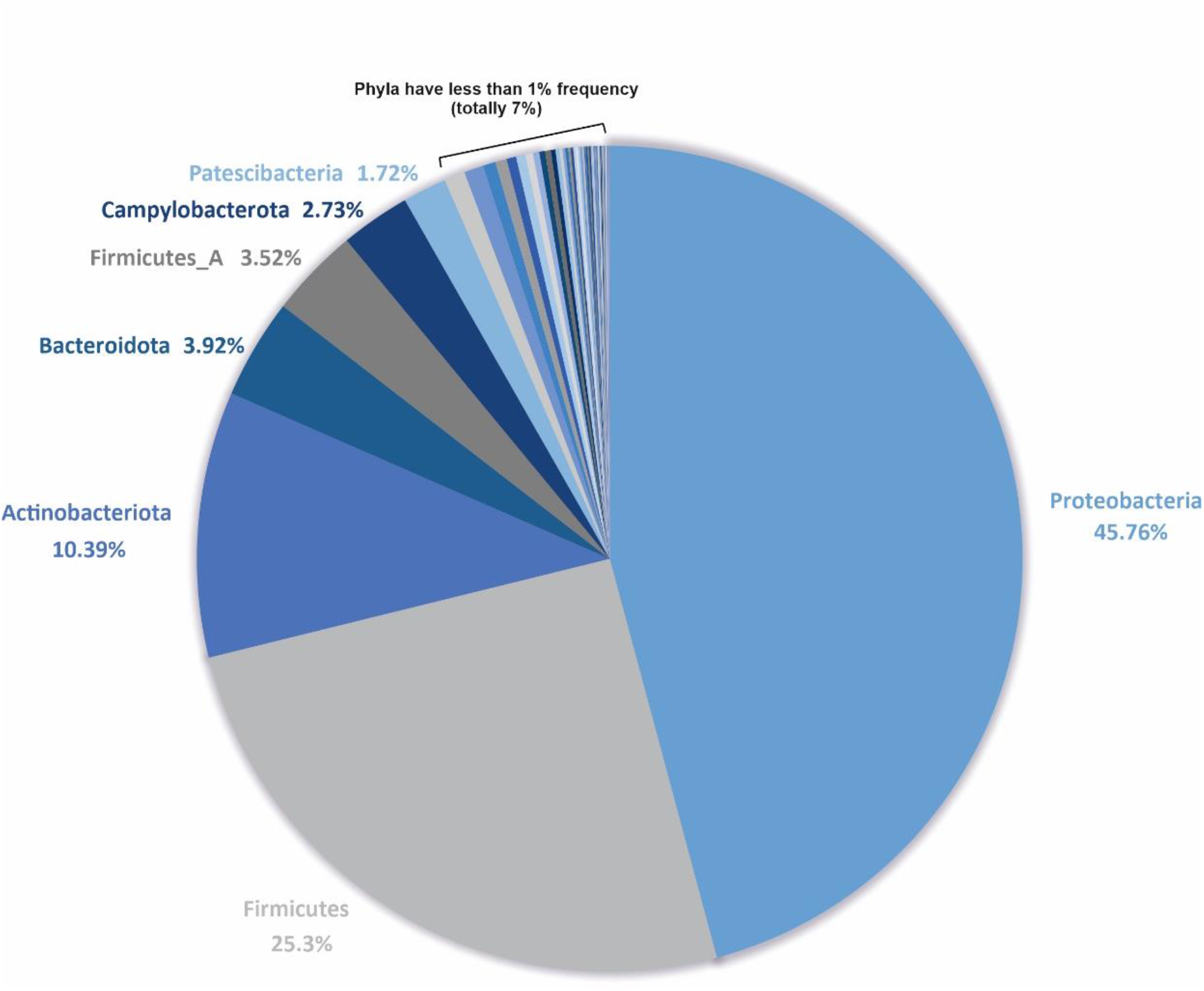
Distribution of 143512 genomes of the GTDB database release 89 in different phyla.

**Supplementary Figure S3.**
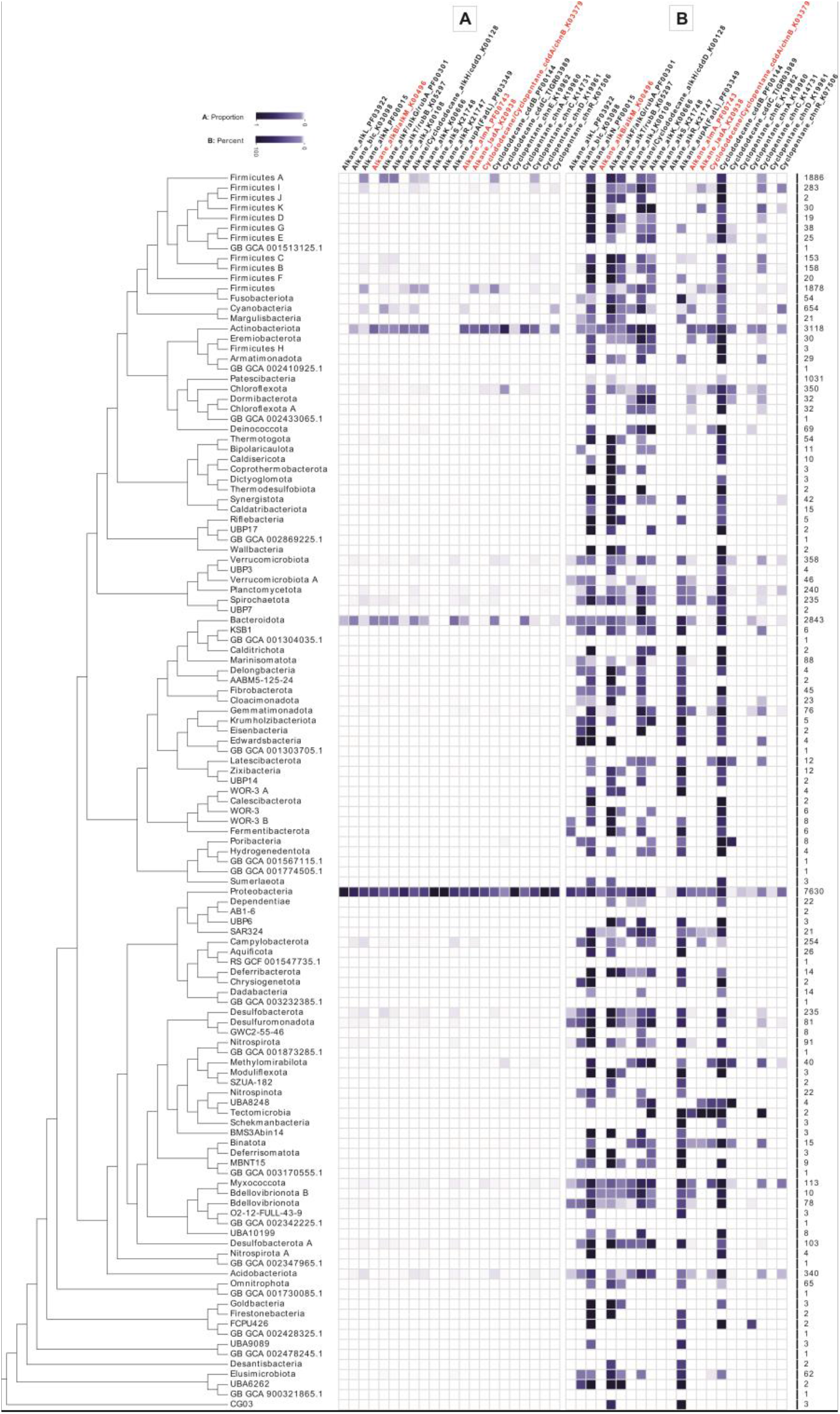
Distribution of aliphatic hydrocarbon-degrading genes across domain bacteria at the phylum level. In plot A, the color gradient indicates the proportion of degrading members of each phylum to the entire HC degrading community. In plot B, the color gradient shows the percentage of HC degrading members of each phylum. Columns are the name of genes involved in HC degradation, which key ones are represented in red.

**Supplementary Figure S4.**
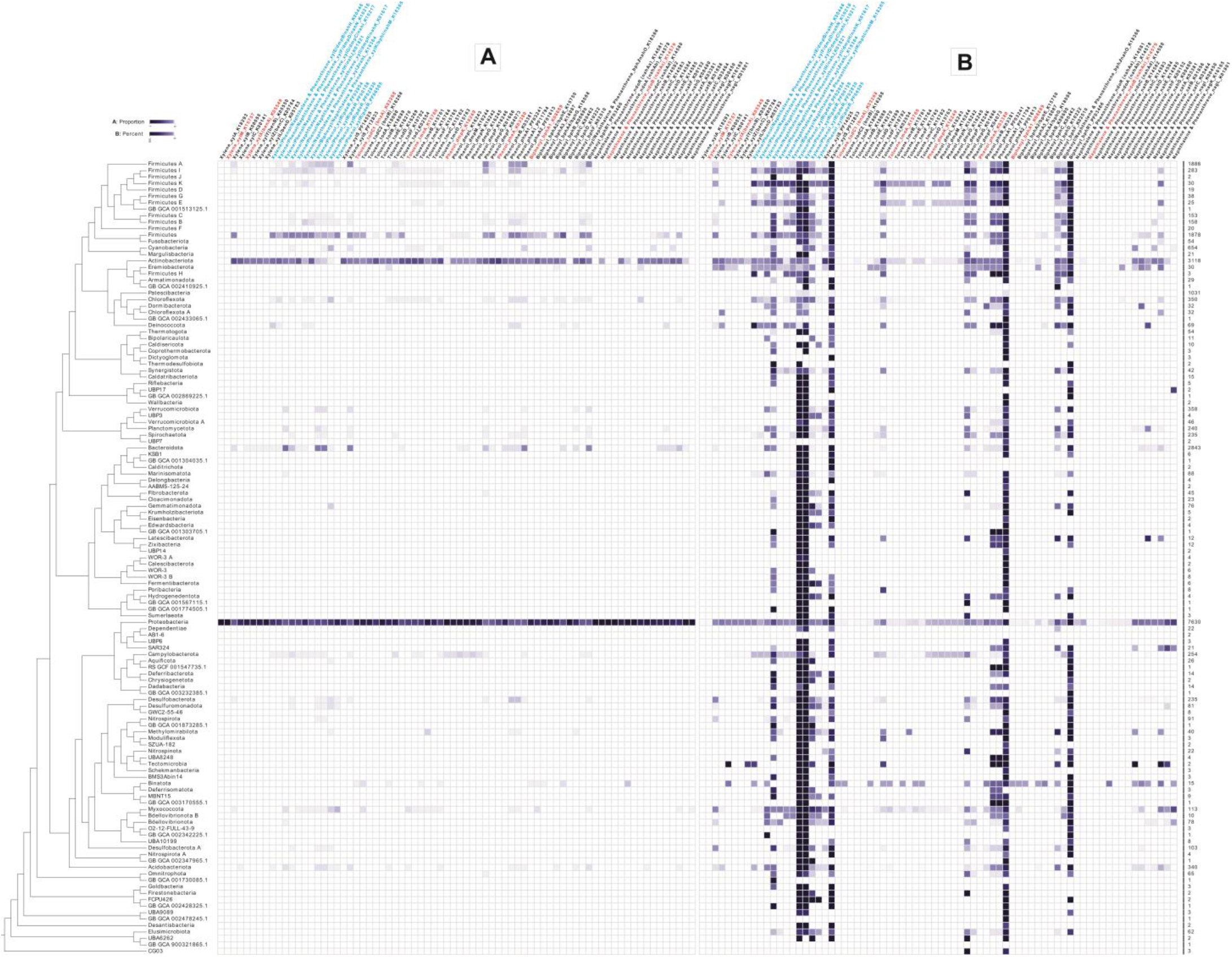
Distribution of aromatic hydrocarbon-degrading genes across domain bacteria at the phylum level. In plot A, the color gradient indicates the proportion of degrading members of each phylum to the entire HC degrading community. In plot B, the color gradient shows the percentage of HC degrading members of each phylum. Columns are the name of genes involved in HC degradation, which key ones are represented in red. Enzymes written in blue are shared among the degradation processes of different aromatic compounds (xylene, phenol and naphthalene).

**Supplementary Figure S5.**
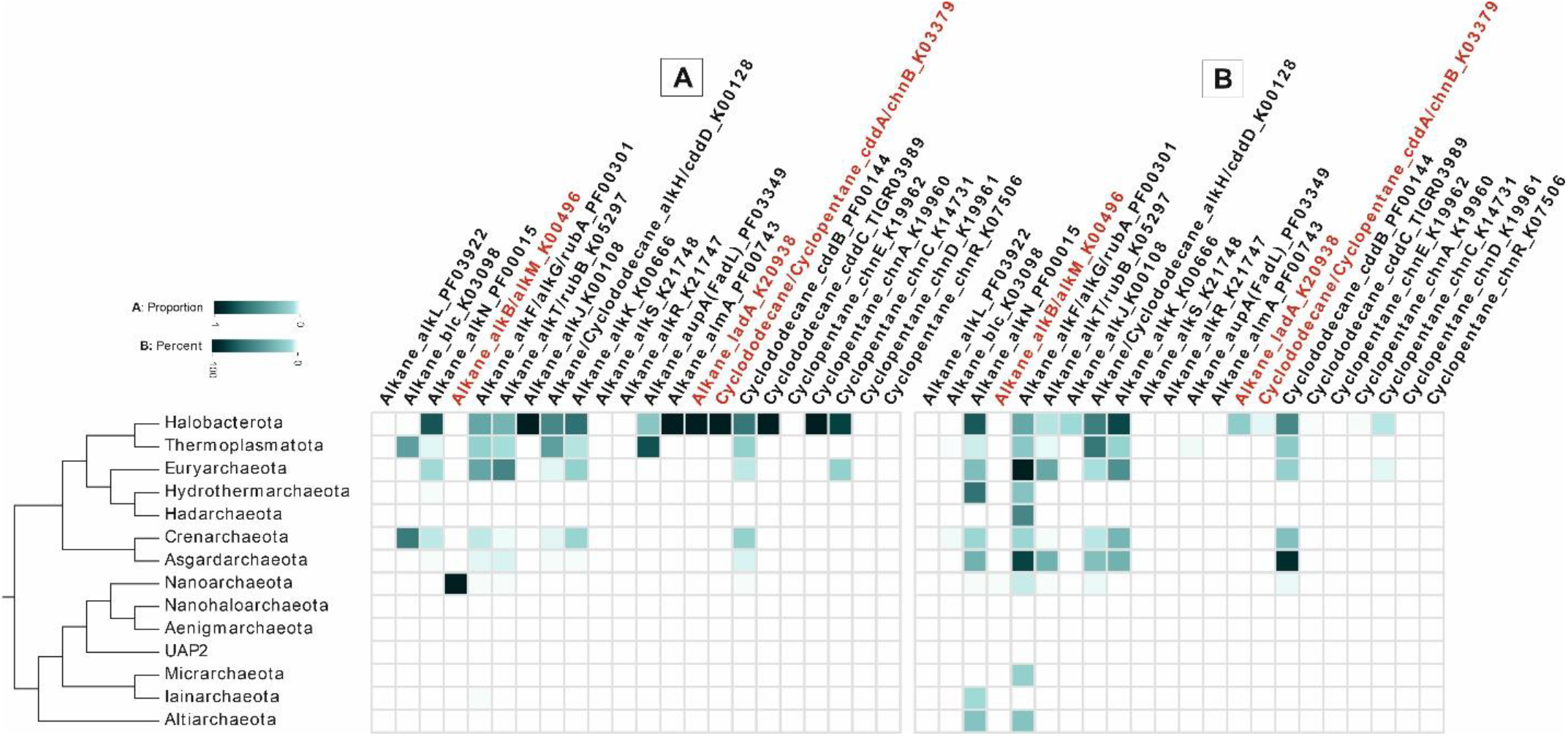
Distribution of aliphatic hydrocarbon-degrading genes across domain archaea at the phylum level. In plot A, the color gradient indicates the proportion of degrading members of each phylum to the entire HC degrading community. In plot B, the color gradient shows the percentage of HC degrading members of each phylum. Columns are the name of genes involved in HC degradation, which key ones are represented in red.

**Supplementary Figure S6.**
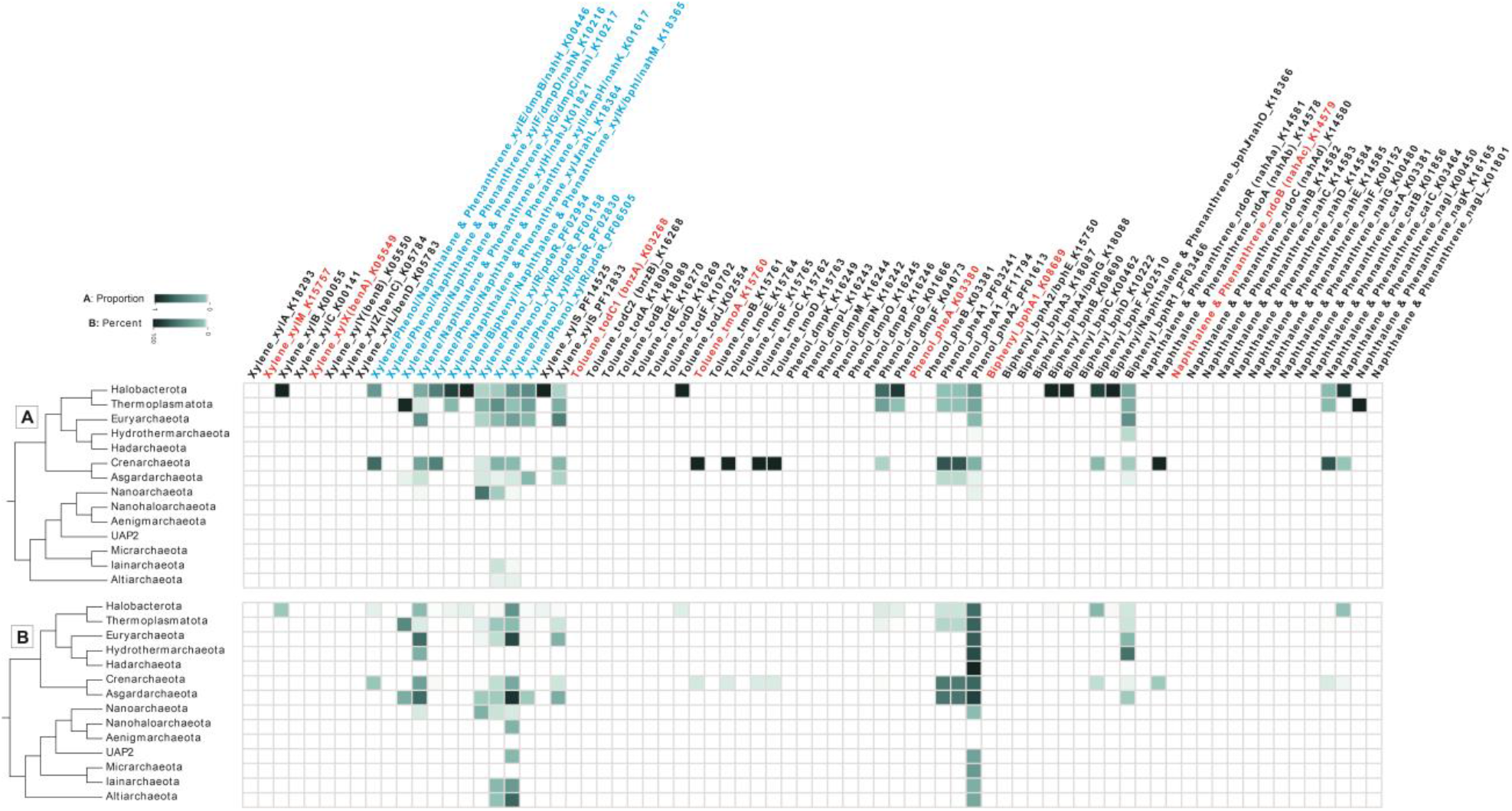
Distribution of aromatic hydrocarbon-degrading genes across domain archaea at the phylum level. In plot A, the color gradient indicates the proportion of degrading members of each phylum to the entire HC degrading community. In plot B, the color gradient shows the percentage of HC degrading members of each phylum. Columns are the name of genes involved in HC degradation, which key ones are represented in red. Enzymes with blue color are shared among the degradation processes of different aromatic compounds (xylene, phenol and naphthalene).

**Supplementary Figure S7.**
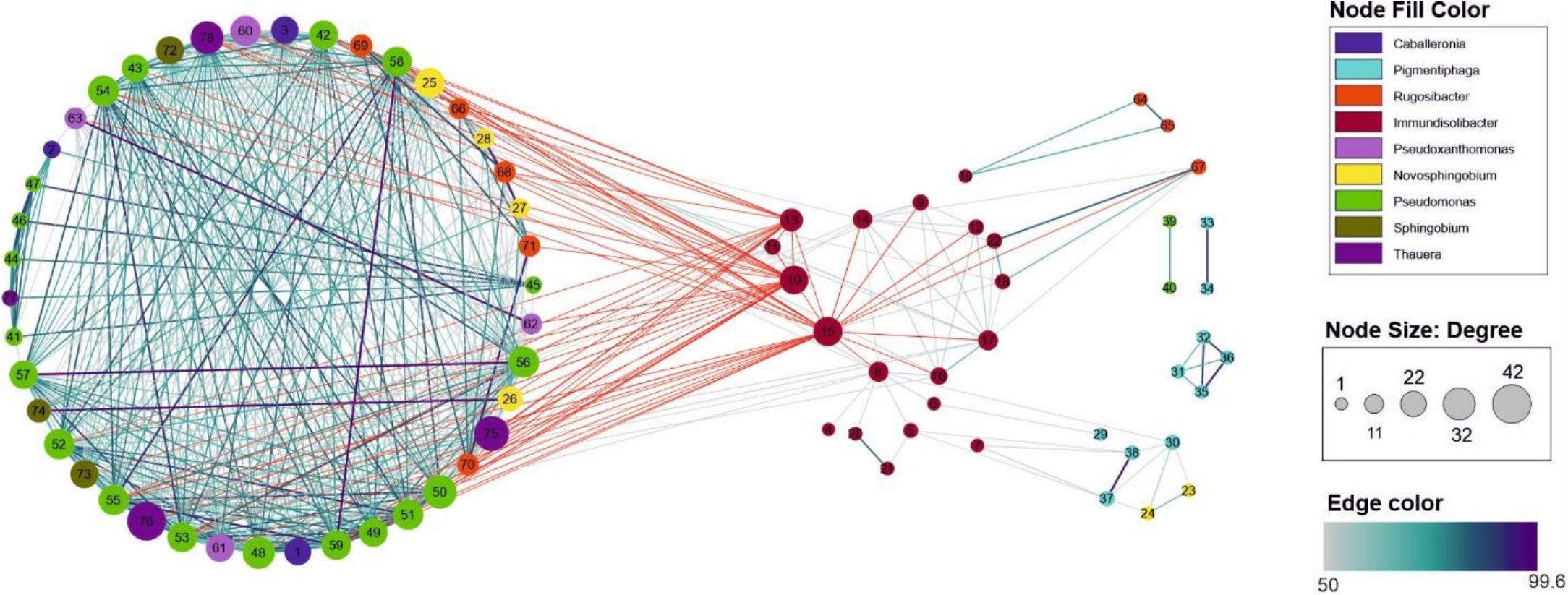
Network interaction between 18 copies of xylX gene in *Immundisolibacter cernigliae* and other genomes with more than two copies of this gene. Only the blast identity values between 50 to 100 percent are shown. Edges are color-coded based on their blast identity. The size of nodes is based on the “Degree,” which is determined by the number of edges of each node. Edges in red are versions of xylX in *Immundisolibacter cernigliae* that had a higher degree than others. The gene ID of the assigned number of each node is represented in Supplementary Table S7.

**Supplementary Figure S8.**
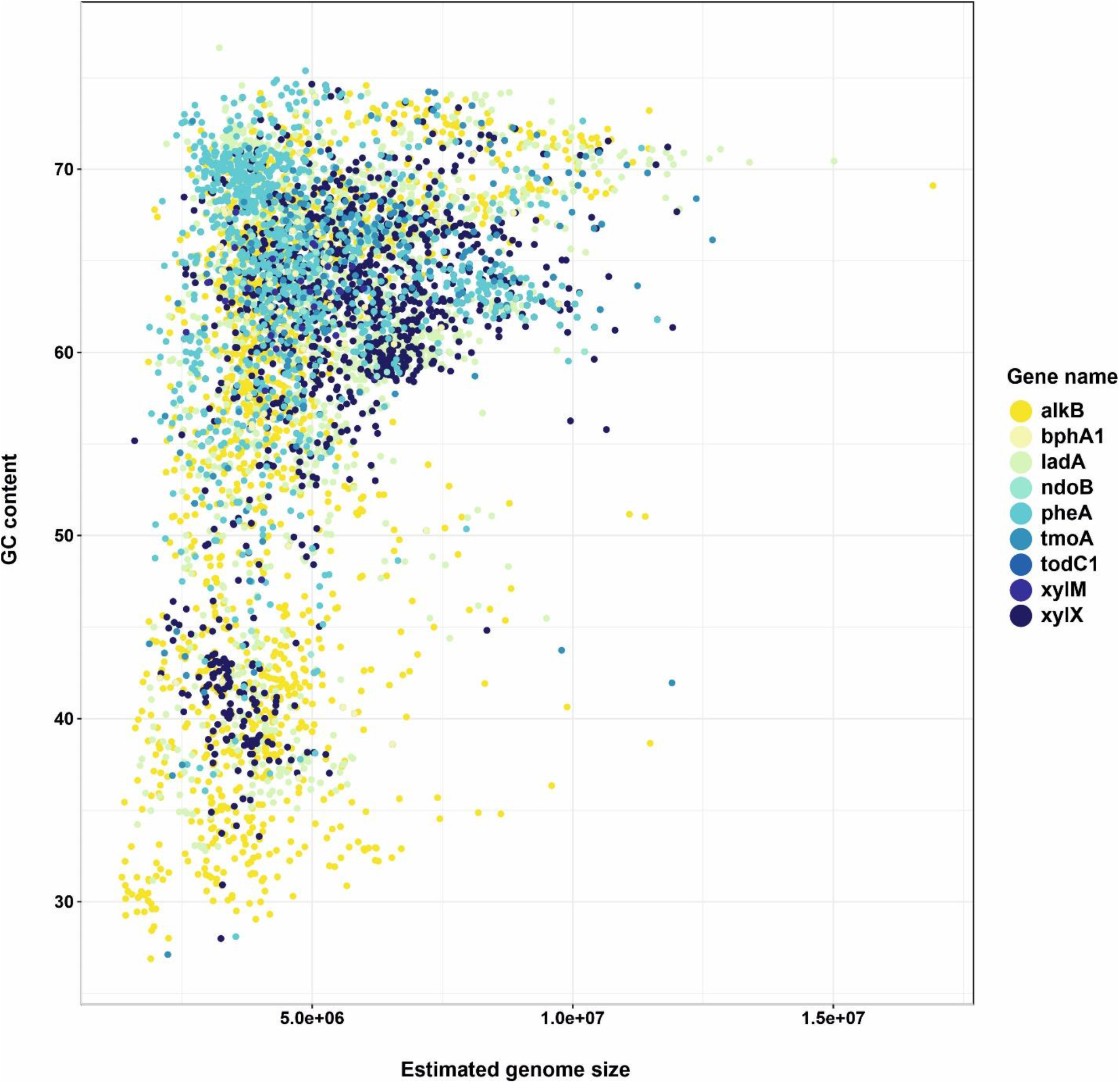
Distribution of genome size versus GC content of the studied genomes with key HC degrading genes.

**Supplementary Figure S9.**
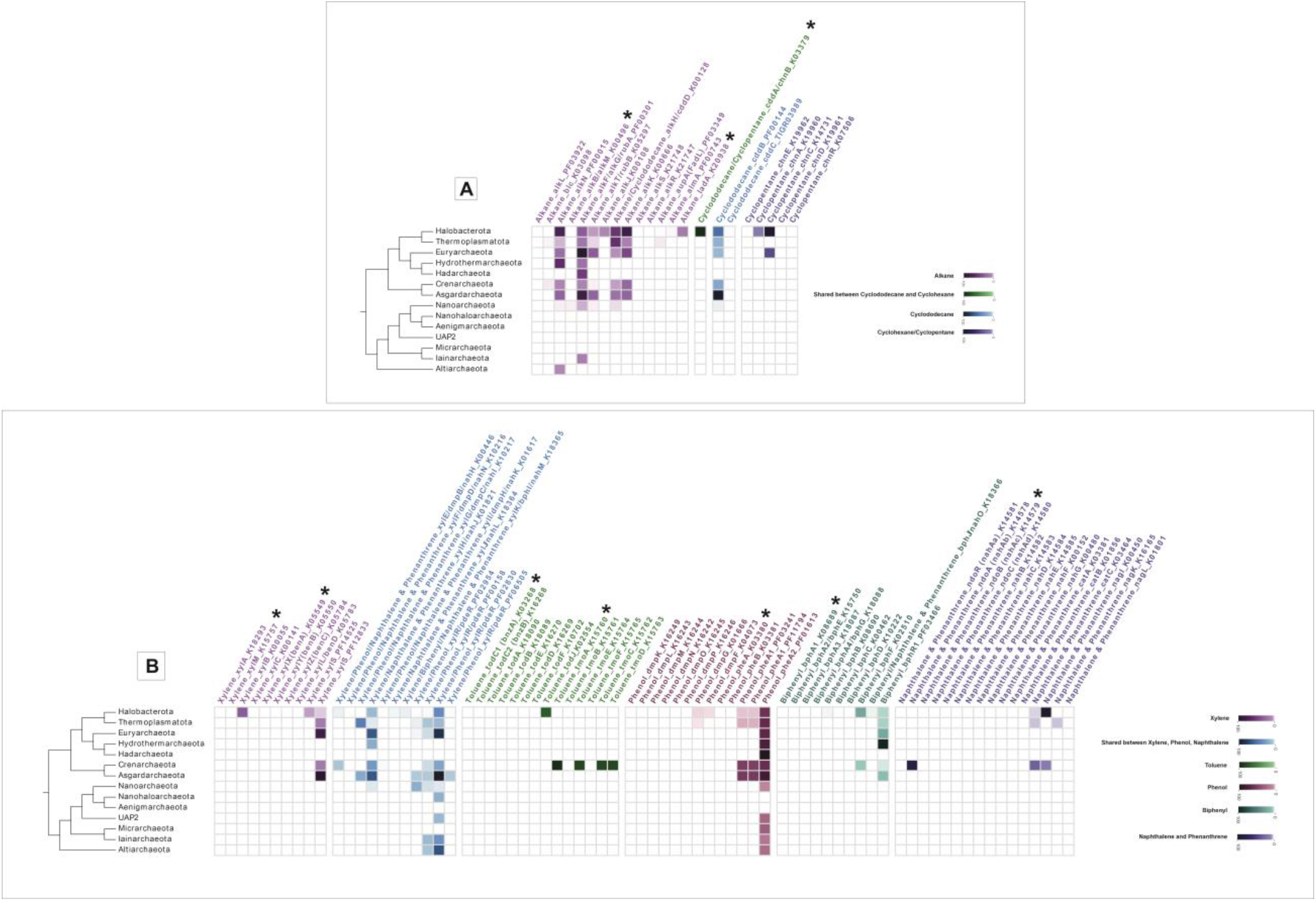
Distribution of aliphatic (A) and aromatic (B) hydrocarbon-degrading genes across domain archaea at the phylum level. Columns show the name of genes involved in HC degradation and are represented in different colors for various compounds. The color gradient for genes of each compound indicates the percentage of HC degrading members of each phylum.

